# The role of oxygen in avascular tumor growth

**DOI:** 10.1101/024562

**Authors:** David Robert Grimes, Pavitra Kannan, Alan McIntyre, Anthony Kavanagh, Abul Siddiky, Simon Wigfield, Adrian Harris, Mike Partridge

**Affiliations:** Cancer Research UK/MRC Oxford Institute for Radiation Oncology, Gray Laboratories, University of Oxford, Old Road Campus, Oxford, OX3 7DQ, United Kingdom; The Weatherall Institute for Molecular Medicine, University of Oxford, John Radcliffe Hospital/Headley Way, Oxford, OX3 9DS, United Kingdom; Advanced Technology Development Group, Department of Oncology, University of Oxford, Old Road Campus Research Building, Oxford, OX3 7DQ, United Kingdom

## Abstract

The oxygen status of a tumor has significant clinical implications for treatment prognosis, with well-oxygenated subvolumes responding markedly better to radiotherapy than poorly supplied regions. Oxygen is essential for tumor growth, yet estimation of local oxygen distribution can be difficult to ascertain *in situ*, due to chaotic patterns of vasculature. It is possible to avoid this confounding influence by using avascular tumor models, such as tumor spheroids, a much better approximation of realistic tumor dynamics than monolayers, where oxygen supply can be described by diffusion alone. Similar to in *situ* tumours, spheroids exhibit an approximately sigmoidal growth curve, often approximated and fitted by logistic and Gompertzian sigmoid functions. These describe the basic rate of growth well, but do not offer an explicitly mechanistic explanation. This work examines the oxygen dynamics of spheroids and demonstrates that this growth can be derived mechanistically with cellular doubling time and oxygen consumption rate (OCR) being key parameters. The model is fitted to growth curves for a range of cell lines and derived values of OCR are validated using clinical measurement. Finally, we illustrate how changes in OCR due to gemcitabine treatment can be directly inferred using this model.

## 1 Introduction

Tumor spheroids are clusters of cancer cells which grow in approximately spherical 3D aggregates. This property makes them a useful experimental model for avascular tumor growth. Spheroids are preferred over 2D monolayers in several applications as the signalling and metabolic profiles are more similar to *in vivo* cells than standard monolayers [1]. Like monolayers, spheroids are relatively straightforward to culture and examine. For these reasons, spheroids have been widely used to investigate the development and consequences of tissue hypoxia. [1]. Early investigations using spheroids began in earnest in the 1970s [2], and the nature of spheroid growth has long been an active question, with several interesting properties mimicing solid tumors. Conger & Ziskin [3] analysed the growth properties of tumor spheroids and noted that they appeared to grow in three distinct stages; exponentially, approximately linearly and then reaching a plateau. A similar type of growth was seen over 15 different tumor cell lines [4], and it was observed that this growth could be approximated to a Gompertzian curve, which described the approximate sigmoidal shape of the growth curves well. In recent years, there has been renewed interest in tumor spheroids in general and the scope for their application has increased dramatically - spheroids have been used in radiation biology [5–8] as a means to test fractionation and other parameters in a controllable environment, in chemotherapy to act as a model for drug delivery [9–12] and even to investigate cancer stem cells [13]. Cancer spheroids have also shown potential as a model for exploring FDG-PET dynamics [14] to explore hypoxia effects in solid tumors.

The distinct sigmoidal growth curves seen in spheroids also occur in some solid tumors, prompting investigation into whether any appropriate sigmoidal curve could be tempered to describe spheroid growth, including the von Bertalanffy and logistic family of models. It has been shown by Feller as early as the 1940s [15] that statistical inference alone could not discriminate between such models; while initially it was postulated that any sigmoid shape may be adequate [16], later analysis [17] found that while the sigmoid shape was a pre-requisite to describe spheroid growth, it is not a solely sufficient condition. Gompertzian models have also been used, and have the advantage of being well suited to situations where empirical models are required, such as the optimization of radiotherapy [17–19]. A hybrid “Gomp-ex” model [20] was also found to fit observed spheroid growth curves well [17]; in this model, initial growth is exponential, followed by a Gompertzian phase when the increasing cell volume reduces the availability of nutrients to tumor cells. While Gompertzian models of growth can describe the growth of tumor spheroids well, they are do not directly address the underlying mechanistic or biophysical processes. Several complex models of avascular growth have arisen from the field of applied mathematics; a review by Roose et al [21] offers an overview of mathematical approaches to modelling avascular tissue, broadly separating published approaching into either continuum mathematical models employing spatial averaging or discrete cellular automata-type computational models. Continuum models are typically intended to model in situ tumors before the onset of angiogensis, and tend to have terms for numerous physical phenomena including acidity and metabolic pathways. Other authors have posited temporal switching of heterogeneous cell types in 2D models in response to a generic growth factor [22], or travelling wave solutions for a model switching between living and dead cell types [23] and even models for the stress cells experience in an avascular tumor [24]. These models include terms for a wide array of intercellular processes with varying levels of mathematical elegance and sophistication, but the presence of a large number of free parameters make direct validation of such models difficult and the models are not always useful or suitable for in *vitro* data. Despite extensive investigation from several avenues, this is still an active problem - a recent review in *Cancer Research* [25] stated that new models and analysis are vital if we are to understand the processes in tumor growth.

In this investigation, we confine our investigation to a simple case to allow us reduce the number of parameters. Specifically, we shall model the effects of oxygen on spheroid growth whilst controlling for other potentially confounding factors. Spheroids provide insight into how avascular tumors propagate; as spheroids increase in size, their central core becomes anoxic and leads to the formation of two distinct zones - a necrotic core and a viable rim, as depicted in fig 1. We have recently derived an explicit analytical model for oxygen distribution in spheroids, which accurately predicts properties such as the extent of the anoxic, hypoxic and viable regions and allows determination of the oxygen consumption rate from first principles for a spheroid at a given time point [26]. It can further be shown from this analysis that the oxygen consumption rate (OCR) of a spheroid drives both its oxygen distribution and the physical extent of the anoxic core *r_n_*. Indeed, oxygen distribution in situ has serious ramifications for therapy too, as radiotherapy is up to a factor of 3 more effective in well-oxygenated regions [27]. In this work, we derive a time-dependent discrete growth model for tumor spheroids, linking their relative rates of oxygen consumption to their growth curves. The model used in this work has only has free parameters of oxygen diffusion and oxygen consumption rate, and average cellular doubling time so that local oxygen partial pressure determines cell behaviour. We do not model other nutrients such as glucose, which are assumed to be present in excess in the media. The model is limited to spheroids from a single cell line. This model can be directly contrasted to experimental data and is validated across a range of cell lines.

**Figure 1.**
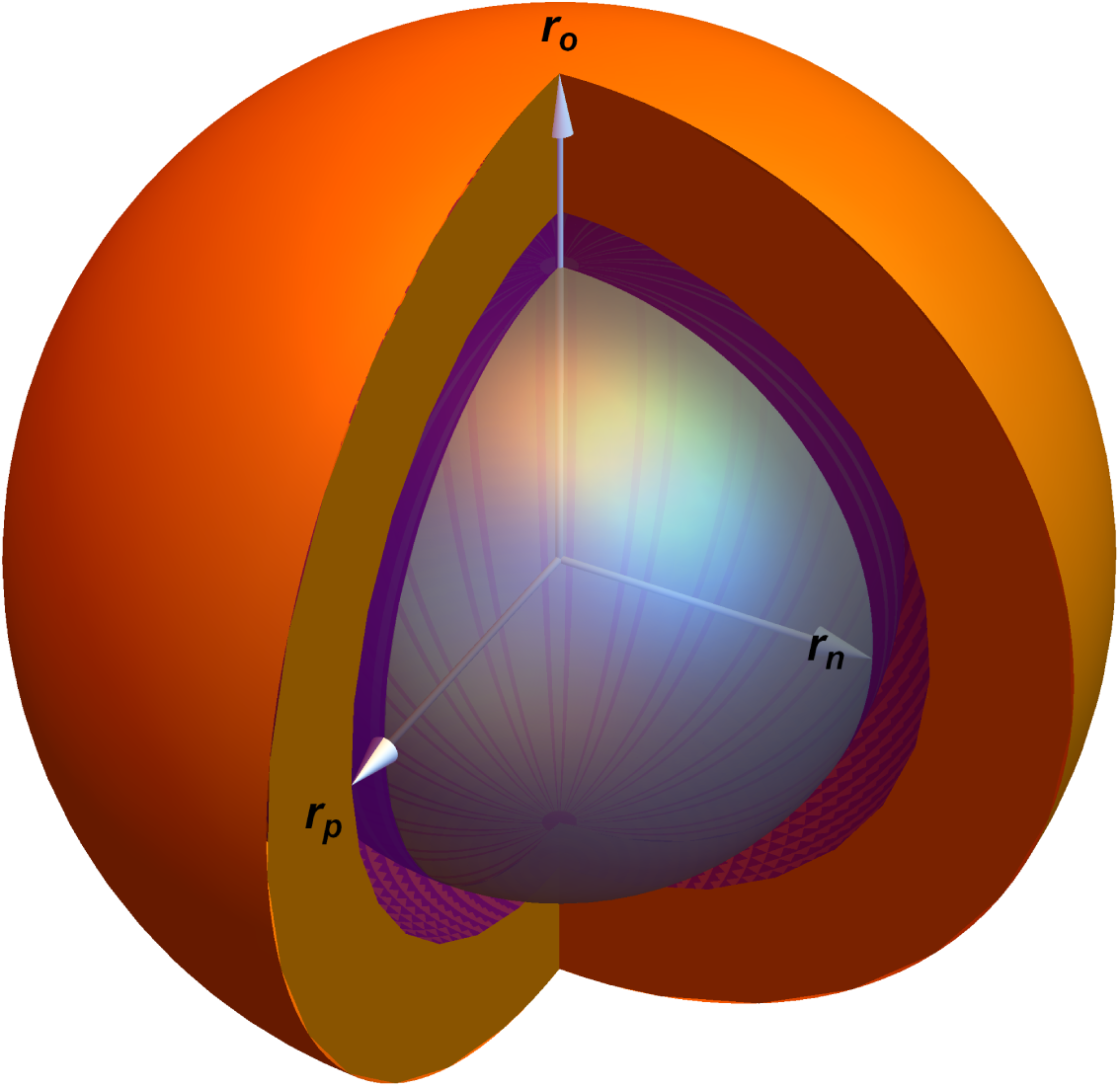
Cross-section of a tumor spheroid of radius *r_o_*. The anoxic radius is denoted by *r_n_*. The radius *r_p_* depicts the radial extent of *p_m_*, the minimal oxygen level required for mitosis. The orange part of the image is the region *r_p_* ≤ *r* ≤ *r_o_*, the purple part corresponds to *r_n_* ≤ *r* ≤ *r_p_* and the central anoxic core (*r* ≤ *r_n_*) is shown in gray.

## 2 Methods

### Oxygen diffusion & derivation of mechanistic growth model

For a tumor spheroid in a medium with external partial pressure *p_o_*, oxygen will diffuse isotropically at a rate *D* throughout the spheroid whilst being consumed by the tumor cells. The rate of oxygen consumption, *a*, has been shown to determine the oxygen tension throughout the spheroid and the resultant boundaries of different spheroid regions [26].

For a spheroid consuming oxygen at a rate *a*, the diffusion length *r_l_* is the maximum radius a spheroid can obtain when the partial pressure at the centre (*r* = 0) is exactly zero. This corresponds to a spheroid with no central anoxia. It can be shown *r_l_* is given by

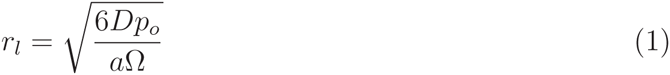

where Ω = 3.0318 × 10^7^ *mmHg kg m*^−3^ is a constant arising from Henry’s law. Oxygen consumption rate (OCR) is usually expressed as volume of oxygen consumed per unit mass per unit time. This can be readily subsumed with the Henry’s law constant to yield *a*Ω, the OCR in units of oxygen pressure per second. For a spheroid with a radius *r_o_* > *r_l_*, a central anoxic core of radius *r_n_* exists, as illustrated in Fig. 1. The anoxic core, *r_n_*, is related to oxygen consumption rate *a* and spheroid radius *r_o_* by

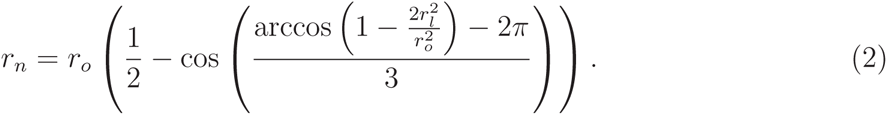

The same analysis [26] allows determination of the partial pressure at any point in the spheroid. If the minimum oxygen tension required for mitosis is *p_m_* where 0 ≤ *p_m_* ≤ *p_o_*, then the radius at which this minimum tension is achieved, *r_p_* is given by

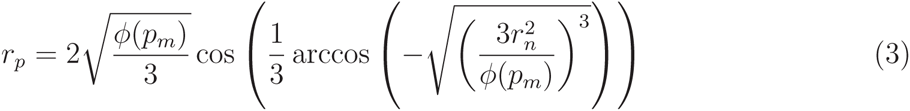

where *ϕ*(*p_m_*) is given by

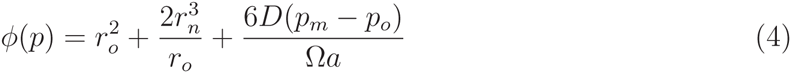

These equations describe the oxygen distribution and physical regions of any given spheroid, but do not state anything about spheroid growth. However, this can be readily extended. Initially we consider the volume of living, viable cells. This is simply the difference in volume between spheres of radius *r_o_* and *r_n_*, expressed as

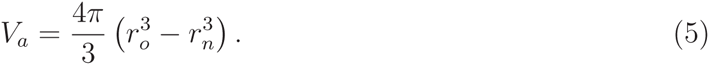

We further assume that cells below a minimal oxygen threshold are unable to undergo mitosis and that *r_p_* is the radius at which *p* = *p_m_*, and *r_n_* ≤ *r_p_* ≤ *r_o_*. The volume of cells able to proliferate is therefore given by

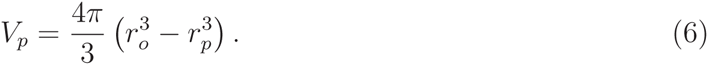

If the average cellular doubling time is *t_d_*, then after this time interval then *V_p_* new cells will be created. For an initially well-oxygenated spheroid with no anoxic core, this increase in volume increases the radius of the spheroid. Increasing spheroid radius changes the oxygen profile as defined by equations 1–4, and an anoxic centre is eventually created. Cells in this domain quickly die and undergo cytolysis, with the spheroid in the medium being readily permeable to resultant water. We further assume that upon creation of the anoxic core, a volume equal to *V_p_* is pushed in towards the central void by a number of mechanisms expanded upon in the discussion section. If the proliferating volume is greater than the anoxic core volume, there is a net increase in spheroid volume and radius. Each increase in spheroid volume results in an increasing anoxic radius, and eventually the volume of proliferating cells is completely contained within the spheroid core, after which no net growth is observed despite there being a radius of proliferating cells. At each doubling time, a changing spheroid volume gives a new radius (*r_oN_*), anoxic radius (*r_nN_*) and proliferating limit radius (*r_pN_* ), all of which can be calculated from oxygen consumption rate a. This can be formulated together as a piecewise iterative model, with total volume and radius at the iteration *N* + 1, given by

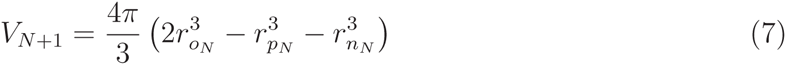

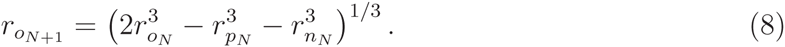

This growth model predicts that for a sufficiently small spheroid (*r* ≪ *r_l_*) that growth is initially exponential, then inhibited by hypoxia and central anoxia. The critical radius at which growth is no longer exponential, and where oxygen consumption limits the proliferating extent occurs at

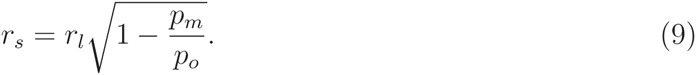

These equations give rise to a classic sigmoid curve, where initial exponential growth is followed by an approximately linear phase before growth begins to plateau. The model predicts that the volume spheroids can obtain and the rate at which they grow is heavily influenced by the OCR and that high consumption rates result in decreased plateau volumes relative to spheroids which consume oxygen at a lower rate. This is an interesting finding, as this models yields a sigmoidal shape akin to the observed experimental curves mechanistically without any *a priori* assumption or forcing - some sample curves are depicted in Figure A in supplementary material **S1**. These equations also predict that oxygen limited growth eventually plateaus, in line with observations by Conger *et al* and others [3] - the radius at which this occurs can be estimated by satisfying

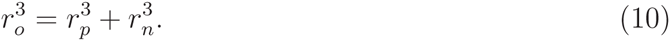

The growth model predicts spheroid growth, using only the parameters of oxygen consumption rate *a* and average cellular doubling time *t_d_*. In this work, we examine the growth curves from spheroids from a number of different cell lines, and also outline a clinical method for estimate OCR so that theoretical fits can be validated to clinical data.

### Measurement of oxygen consumption and relationship with cellular mass

There is a degree of ambiguity in clinical terminology that is worth addressing here - the quantity of oxygen gas consumed per unit time can be measured using extracellular flux methods. However, this is a relative measurement of oxygen consumed per cell per unit time. As a consequence, this could change markedly between cell lines where average cell mass is different. In this work, we define OCR as oxygen consumed per unit time per unit mass in line with previous work [26, 28]. This has the advantage of facilitating cross-comparison of oxygen consumption between cell lines. Extracellular flux methods for calculating oxygen consumption rate per cell works by isolating an extremely small volume (typically less than 7*μ*l) of medium above a monolayer and measuring changes in the concentrations of dissolved oxygen for a number of cells *N_c_*. If we denote measured extracellular flux values as *S_H_* (in units of moles of oxygen per cell per minute), then it is relatively simple to convert this to general OCR *a* (in S.I units of m^3^ kg^−1^ s^−1^ ). This requires an estimate of average cellular mass for a given cell lines, *m_c_*. The full conversion is given by

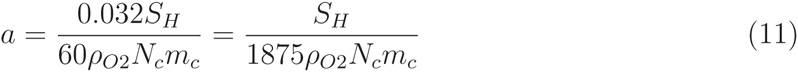

where *ρ*_*O*2_ is the density of oxygen gas (taken to be *ρ*_*O*2_ = 1-331 kg m^−3^) and *N_c_* is the number of cells in the sample. The factor of 0.032 in the numerator arises when converting from moles to kilograms, given the fact that a mole of oxygen gas has a mass of 32g [26]. The factor of 60 in the denominator converts from minutes to seconds. Equation 11 indicates that *a* ∝ *S_H_* and 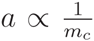, so information about cellular mass is needed to completely describe the oxygen consumption characteristics of a cell line. If cell volume can be estimated, then cell mass may be inferred by assuming that cells have the density of water *ρH*2*O*. Mass is then given by

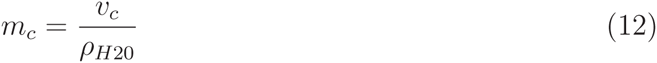

Oxygen consumption rate was measured using a Sea-horse extracellular flux analyzer (Seahorse Bioscience, Massachusetts) and a range of values of *S_H_* for a range of cell numbers to ensure linearity for each cell line. The raw Sea-horse data is tabulated in Tables A-F inclusive in supplementary material **S1**.

### Mass estimation technique

As cellular mass *m_c_* is related to the oxygen consumption rate by equation 5, it was important to estimate this to facilitate comparison between OCR derived from theory and experiment. To obtain approximate values for cell mass, we estimated the volume of individual cells using 3D confocal microscopy. Cells (1 × 10^5^ per well) from all lines were seeded onto glass coverslips in a 6-well plate and incubated at 37°C for 24 hours; all subsequent experiments were carried out at room temperature. Once cells had attached to the coverslip, they were washed in serum-free culture medium and then incubated for 5 min with PKH26 (2 *μ*M; Sigma-Aldrich) prepared in Diluent C (Sigma-Aldrich) to fluorescently label the cell membrane. An equal volume of 100% fetal bovine serum was added for 1 min to stop the reaction. Cells were washed with regular culture medium, fixed for 10 min with 4% paraformaldehyde, washed with phosphate-buffered saline, and incubated with Hoechst 33342 (3.2 *μ*Μ; Life Technologies) for 10 min to label nuclei. Coverslips were then washed in PBS and mounted using *SlowFade*^®^ Gold Antifade Reagent (Life Technologies) onto glass slides. Five fields of view from each cell line were acquired using the 20x/0.87 M27 objective on a Zeiss LSM 710 microscope (Carl Zeiss AG). To obtain cell volume estimates, each field of view was scanned using the z-stack feature, with a 1 *μ*m slice thickness, to obtain images for cell volume estimates as shown in Fig. 2.

**Figure 2:**
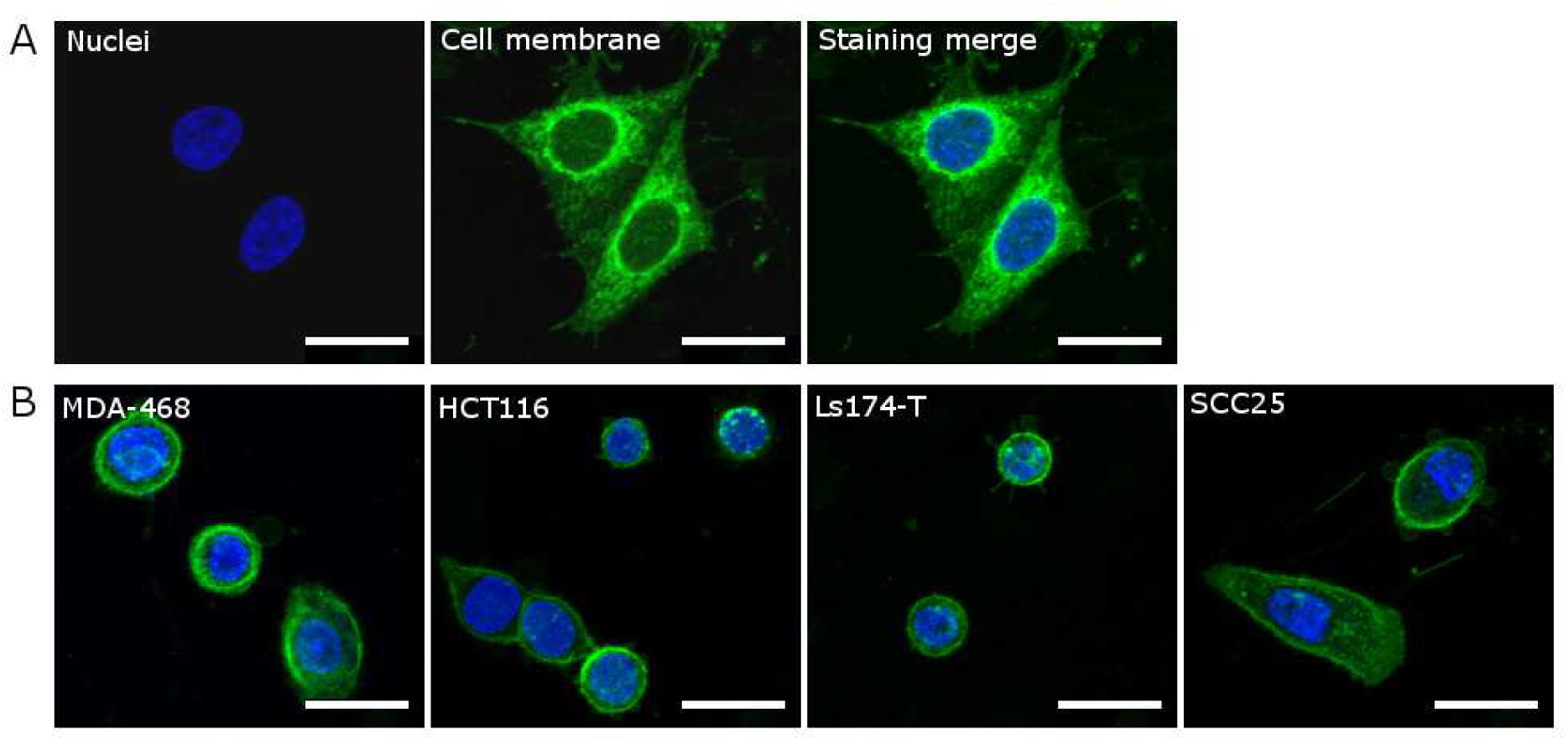
Representative images (maximum intensity projections) of five cell lines that were fluorescently labelled for estimating cell volume using 3D confocal microscopy. Merged images show the simultaneous staining of the nucleus (blue, Hoechst 33342, 3.2*μ*Μ) and the cell membrane (green, PKH26, 2*μ*Μ) in (A) HeLa cells and in (B) the MDA-468, HCT116, Ls-174T, and SCC25 cells. Scale bars represent 10*μ*m

To help counteract inherent blurring, unsharp masking was performed using a blurring kernel and magnitude set manually to provide maximum edge definition. After the unsharp masking technique was applied, a MATLAB script was run which calculated the area of each slice through a cell. These areas were summed and multiplied by the slice thickness (1 *μ*m) to estimate cell volume. Cell mass was estimated using the density transform in equation 12. This method was validated by performing it on Hela cells, and comparing the results to literature estimates of HeLa mass [29].

### Cell culture and spheroid growth

Seven cell lines (ATCC) from different human cancers were cultured for in vitro experiments: the cervical carcinoma line HeLa (ATCC^®^ CCL-2), colorectal lines HCT 116 (ATCC CL-247), LS 174T(ATCC CL-188), the breast cancer lines MDA-MB-468 (ATCC^®^ HTB-132) and MDA-MB-231 (ATCC^®^ HTB-26), the squamous cell carcinoma line SCC-25 (ATCC CRL-1628), the glioblastoma cell line U-87 MG (ATCC HTB-14). All cell lines were grown in Dulbecco’s Modified Eagle Medium (Life Technologies) supplemented with 10% fetal bovine serum, with the exception of the SCC-25 line, which was cultured in growth medium recommended by the supplier (American Type Culture Collection). Cells were maintained at 37°C in humidified incubator containing 5% CO_2_.

Spheroids were generated as previously described [30] in literature. Briefly, 0.2 *μ*l of 2.5 ×10^4^/ml cell suspension was added to each well of a 96-round-bottom-well ultra low attachment plate (Corning Incorporated) in a high glucose medium (4.5g /L). For MDA-MB-468 and SCC25 cells, Matrigel (BD Bioscience) was added at a final concentration of 5%. Plates were then centrifuged at 300×g for 10 min. Centrifugation was carried out at 4°C if Matrigel was required for spheroid formation. Spheroids were maintained at 37°C in a humidified incubator containing 5% CO_2_. Pictures of spheroids were taken using the EVOS XL Core Cell Imaging System (Life Technologies) and volumes were analysed using the ImageJ (NIH). Hela cells were not grown as spheroids, but were used to validate the mass estimation method outlined previously. The oxygen diffusion constant *D* was taken to be close to that of water, approximately *D* = 2 × 10^−9^ m^2^/s [14,26,28]. To minimise fluctuations in glucose concentration, we renewed spheroid medium every 48 hours by replacing half the volume of medium in each well with the equivalent volume of fresh culture medium.

### Model validation

Growth curves were obtained for spheroids for each cell line — table 1 lists a summary of all cell-lines. The initial volume of each spheroid line was measured and from this *r_o_* calculated. The oxygen limit for mitotic arrest was taken from literature to be *p_m_* = 0.5 mmHg for spheroids in a glucose solution [31]. The best-fit between model and the experimental growth curves was then calculated. When possible, the OCR for each cell line was then estimated using the experimental procedure outlined. This data was then used to produce a theoretical growth curve which could be directly compared to the experimental data without degenerate fitting.

**Table 1:** Summary of Spheroids used for growth curves

| Cell Line | Spheroid Formation Method | Number of spheroids |
| --- | --- | --- |
| MDA-MD-231 | Suspension in low-attachment plate | 18 |
| U-87 | Suspension in low-attachment plate | 18 |
| SCC-25 | Suspension with 5% Matrigel in low-attachment plate | 18 |
| HCT 116 | Suspension in low-attachment plate | 24 |
| LS 174T | Suspension in low-attachment plate | 30 |
| MDA-MB-468 | Suspension with 5% Matrigel in low-attachment plate | 18 |

The model could also be used to estimate the effect of different clinical compounds of OCR - to investigate this, 11 HCT 116 spheroids were grown, and 7 of these treated with 50nM of gemacitabine. The remaining 4 functioned as experimental controls. These were stained with the proliferation marker Ki-67 and the hypoxia marker EF5, and sectioned through the centre. The OCR was estimated used sectioning techniques [26] and the resulting OCRs were compared for both groups using a two-tailed Welch’s correction t-test.

## 3 Results

### Model fitting

Theoretically derived growth curves were fit to experimental data for a range of cells lines, and from the best fit parameters used to estimate OCR *a* and average cellular doubling time *t_d_*. This analysis imposes a constraint condition on the data of

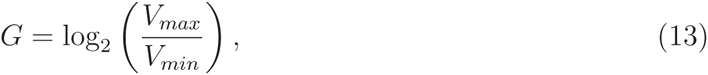

where *V_max_* is the maximum spheroid volume at the end of the growth period and *V_min_* the initial volume at *t* = 0. For model fitting purposes, the condition *G* ≥ 2 ensures that a least two cell doubling times of *t_d_* have transpired and avoids over-fitting. Such a consideration ruled out the use of certain cell lines such as T-47D (ductal carcinoma) as volume increases were too small for analysis over the growth period. Growth curves were fitted for three distinct cell lines; MDA-MB-231 Breast adenocarcinoma, U-87 Glioblastoma, and SCC-25 squamous cell carcinoma. Fits were also obtained on previously published data by Freyer [4, 17] for V-79 hamster fibroblast cells. This data was selected as it is relatively long range ( 60 days) and plateau effects can be readily observed. Fig. 3 shows the ideal theoretical fits for these curves which yields the greatest co-efficient of determination, indicating that the model fits the data extremely well. Error bars on the time axis are 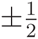 a day to capture uncertainty on exact time which growth curves were measured on a daily basis. It is important to note however that doubling time *t_d_* and diffusion limit are degenerate parameters and there exist a considerable range of parameters which will yield similar fits. This degeneracy is explored further in the discussion.

**Figure 3:**
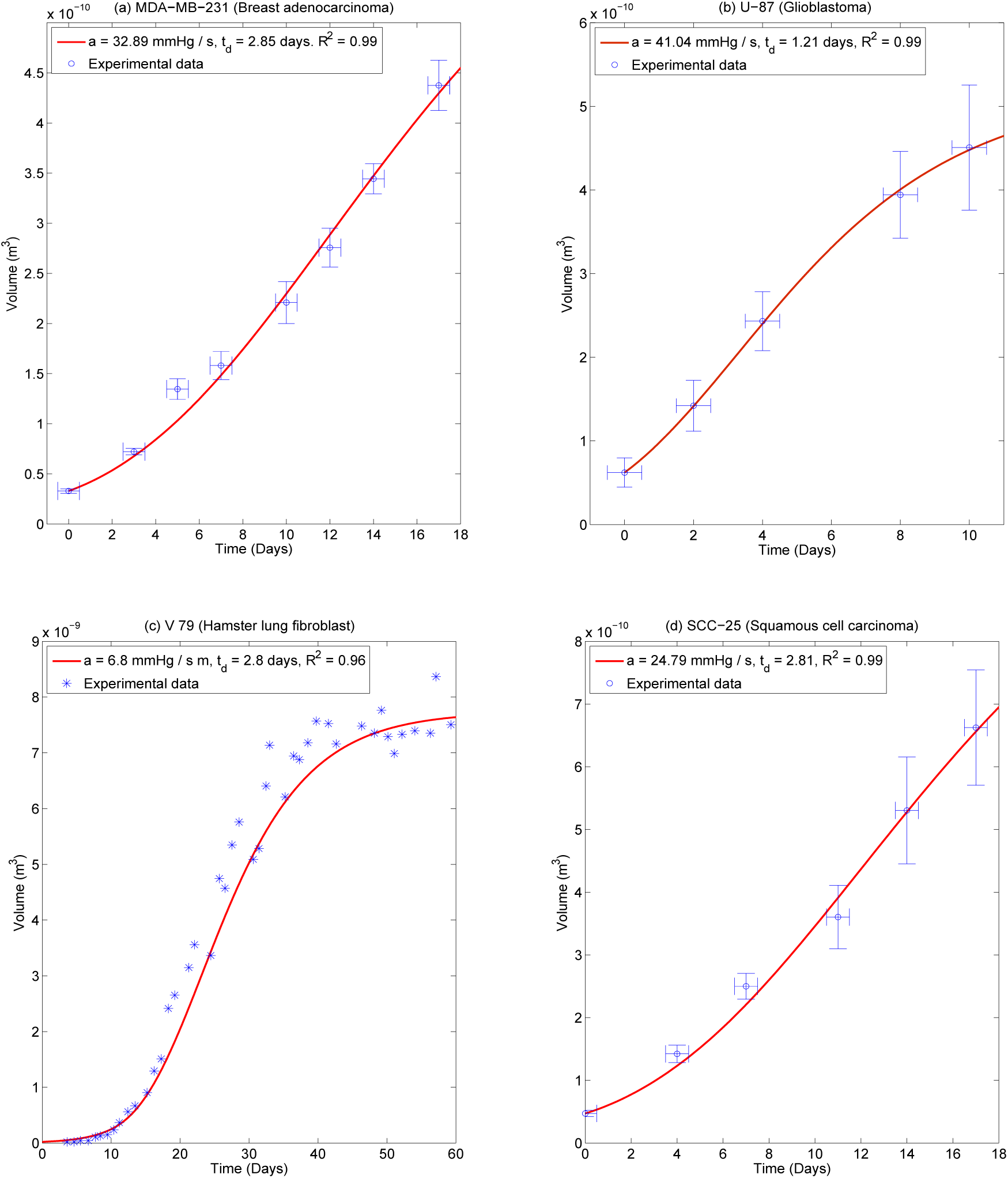
Theoretical best fits for (a) MDA-MB-231 (b) U-87 and (c) Hamster V-79 spheroids. Data for the V-79 cells is from previously published investigations by Freyer [4] and standard errors are not shown on this plot. (d) SCC-25. While best fits are shown in this figure, there are several possible combinations of diffusion limit (*r_l_*) and doubling time (*t_d_*) that produce similarly high co-efficients of determination so these results may not be uniquely determined.

### Comparison of theoretical curves to experimental data

As curve-fitting suggests the model fits the data well, it is possible for some cell lines to avoid potential degeneracy and directly contrast theoretical curves with experimental data, provided OCR can be determined. In this case curve-fitting is not required and model and data can be directly compared. Cell volume and hence mass estimates were obtained for a number of cells in four distinct cell lines; HCT 116 (n = 36), LS 147T (n = 36), MDA-MB-468 (n = 27) and SCC-25 (n = 22). For these cell lines, multiple individual cells could be isolated and cell mass was estimated by the procedure outlined in the methods section. This was combined with the extracellular flux measurements *S_H_* to yield an estimate for the consumption rate *a* and the resultant OCR *a*. Best estimates for oxygen consumption are shown in table 2. The diffusion constant was assumed to be close to that of water so *D* = 2 × 10^−9^ m^2^ s^−1^. The oxygen partial pressure in the medium at the spheroid boundary was *p_o_* = 100 mmHg. Results for these cell lines are shown in fig 4, where model results are contrasted to experimental data. Results are shown with their respective best fit doubling times, *t_d_*. Error bars on the time axis are 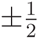 a day to capture uncertainty on exact time which growth curves were measured on a daily basis. The model data illustrated in fig 4 are independent of fitting, directly contrasting the model with OCR taken from the experimental data in table 1.

**Table 2:** Experimentally measured cell mass / OCRE

| Cell Line | Mass (ng) | Seahorse data (pM / cell) | $a \text{ (m}^3\text{kg}^{-1}\text{s}^{-1}\text{)}$ | $a \text{ (mmHg / s)}$ |
| --- | --- | --- | --- | --- |
| HCT 116 | $2.34 \pm 0.45$ | $5.37 \pm 0.13 \times 10^{-3}$ | $9.21 \pm 1.99 \times 10^{-7}$ | $27.92 \pm 6.05$ |
| LS 174T | $2.12 \pm 0.39$ | $3.59 \pm 0.10 \times 10^{-3}$ | $6.80 \pm 1.45 \times 10^{-7}$ | $20.61 \pm 4.38$ |
| MDA-MB-468 | $3.22 \pm 0.61$ | $4.79 \pm 0.30 \times 10^{-3}$ | $5.97 \pm 1.49 \times 10^{-7}$ | $18.08 \pm 4.53$ |
| SCC-25 | $3.54 \pm 1.06$ | $3.27 \pm 0.37 \times 10^{-3}$ | $3.70 \pm 1.53 \times 10^{-7}$ | $11.21 \pm 4.63$ |

**Figure 4:**
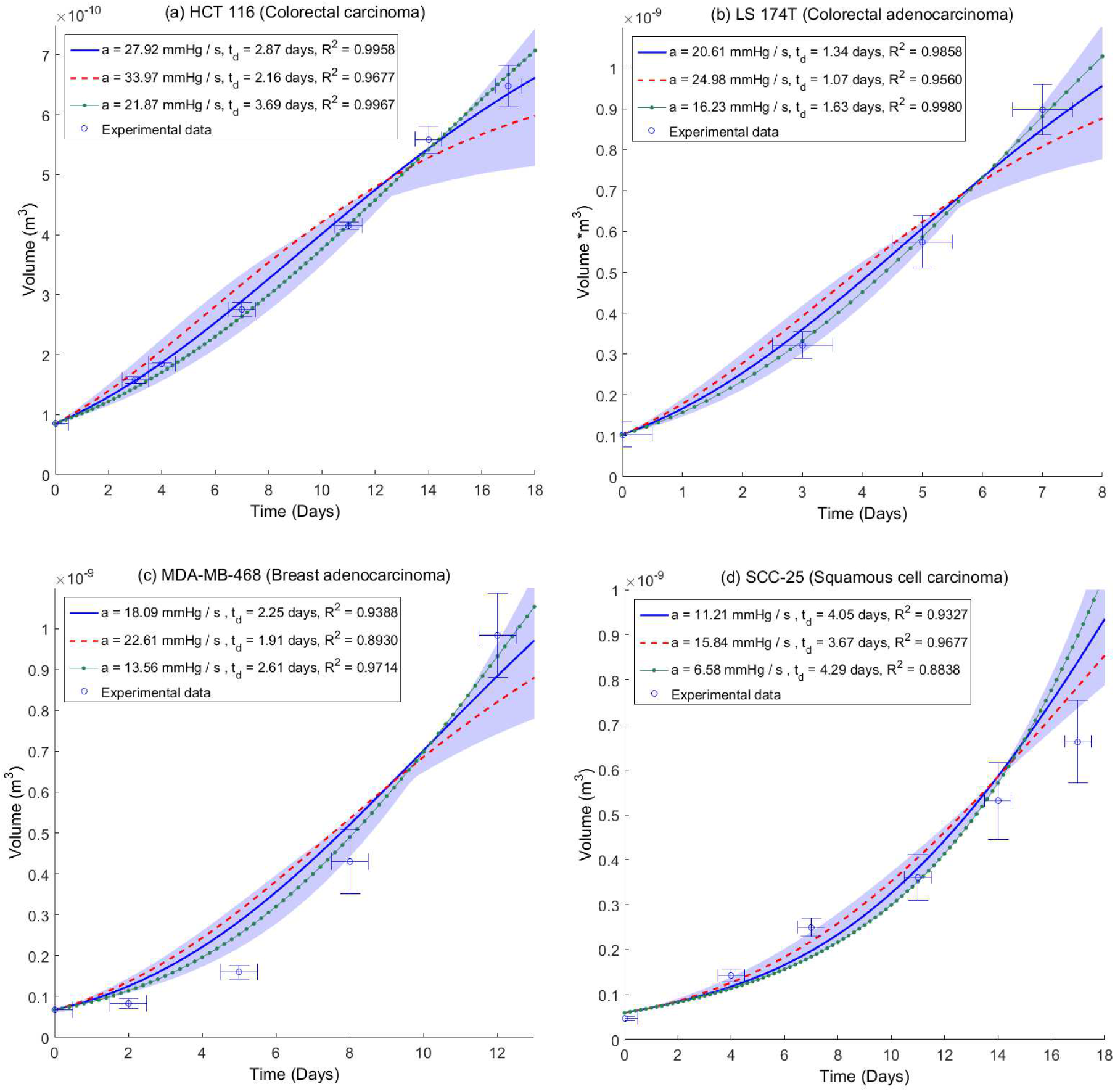
Plots of experimental data and model growth curves for (a) HCT 116 (b) LS 174T (c) MDA-MB-468 and (d) SCC-25 spheroids. In all plots the growth curve due to mean experimentally estimated OCR a is denoted by a solid blue line, with one standard deviation above average OCR marked by a dashed red line and one standard deviation below average consumption marked with a dotted green line. Best fit doubling times *t_d_* and co-efficient of determination are shown for each value with high goodness of fit obtained for each estimated consumption rate within the confidence intervals of experimental data. The shaded area corresponds to range of ± 2 standard deviations for OCR.

### Effects of clinical compounds on OCR

Using the model, the effects of gemcitabine on OCR could also be ascertained. This is illustrated in fig 5, for untreated HCT-116 spheroids and HCT-116 spheroids treated with 50nM of gemcitabine. Untreated HCT 116 spheroids were estimated to have an average OCR of 27.43 mmHg/s, in high agreement with estimated consumption rate experimentally derived in this work (mean value 27.92 mmHg/s) through the confocal mass estimation method. A Welch’s correction two-tailed T-test was performed between the two groups, with highly significant result of *P* < 0.01. These results are shown in Fig. 5.

**Figure 5:**
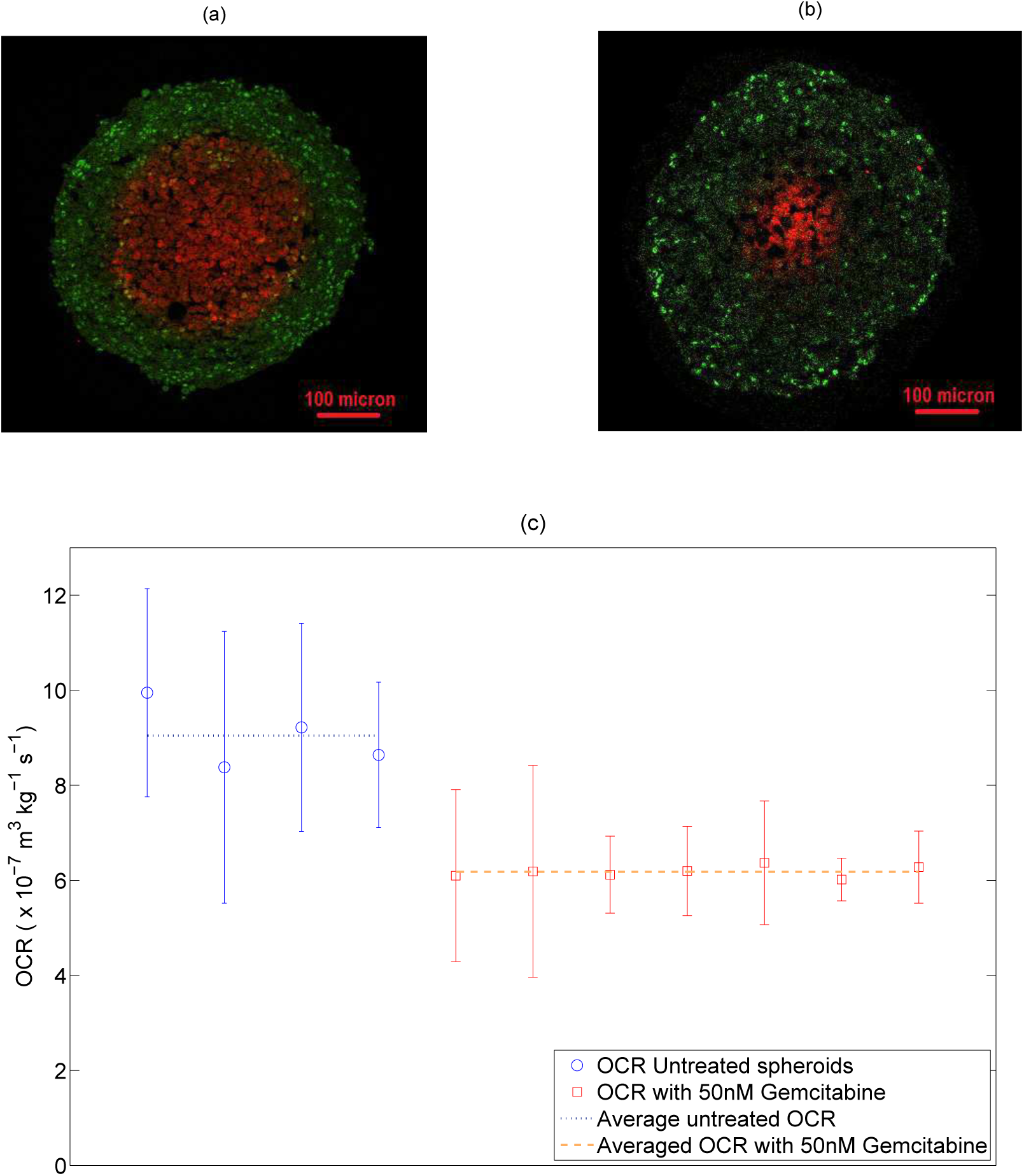
(a) A HCT 116 control spheroid stained for proliferating cells using Ki-67 (green) and for hypoxia using EF5 (red) (b) a HCT 116 spheroid treated with 50 nM of gemcitabine showing markedly smaller hypoxic centre than untreated spheroid. (c) OCR estimated from stained cross-sections by previously outlined method [26] for 4 control spheroids and 7 spheroids treated with 50nM gemcitabine. Average OCR for treated spheroids is 6.18 ×10^−7^ m^3^ kg^−1^ s^−1^ (18.75 mmHg / s) versus 9.05 ×10^−7^ m^3^ kg^−1^ s^−1^ (27.43 mmHg / s) for untreated spheroids (P-Value < 0.01 using a two-tailed Welch’s correction t-test,α = 0.05). This suggests a marked decrease in OCR for treated spheroids.

### Mass validation

To quantify the accuracy of the image analysis method outlined in the prior section, HeLa cells were imaged with the confocal microscope and subsequently deblurred using the unsharp masking technique and area detection algorithm. A selection of these cells (n = 15) were then run through the area detection code in MATLAB so that volume, and hence mass, could be estimated. This yielded an estimated mass of 2.95 ± 0.54 ng for the HeLa cells analysed. This is in good agreement with mass estimates using a cantilever method [29] (3.29 ± 1.14 ng).

## 4 Discussion

This work outlines a simple discrete model for spheroid and avascular tumor growth, quantifying proliferating volume after mitosis with OCR and doubling time as the free parameters. The simple model outlined in this work includes only a minimum of terms for which parameters are either measurable or known. Despite its relative simplicity, the model replicates the growth behaviour of spheroids well for a range of different cell lines of different types, providing a mechanistic explanation for the observed sigmoidal curves associated with spheroid growth. What is particulary worthy of note is the effect of consumption rate on the growth that this analysis suggests, with higher rates of oxygen consumption resulting in a markedly lower plateau volumes. The relatively intuitive reason for this is that the rate of oxygen consumption directly influences the size of the viable spheroid rim [26], as the distance oxygen diffuses is related to how rapidly the respiring tissue consumes it. Consequently, increased oxygen consumption suggests a greater anoxic centre and thinner viable rim, and a smaller volume of proliferating cells.

For the MDA-MB-231, U-87, SCC-25 and V-79 cell lines, theoretical fitting methods were employed to find best OCR and doubling time parameters for a given growth curve. The resultant fits in good agreement (0.96 < *R*^2^ < 0.99) for all cases. There is some unavoidable uncertainty on these fits due to the fact that doubling time *t_d_* and diffusion limit *r_l_* (and by extension OCR) are degenerate parameters, as illustrated in Fig. 6. In principle if the OCR can be estimated, then this degeneracy can be circumvented.

**Figure 6:**
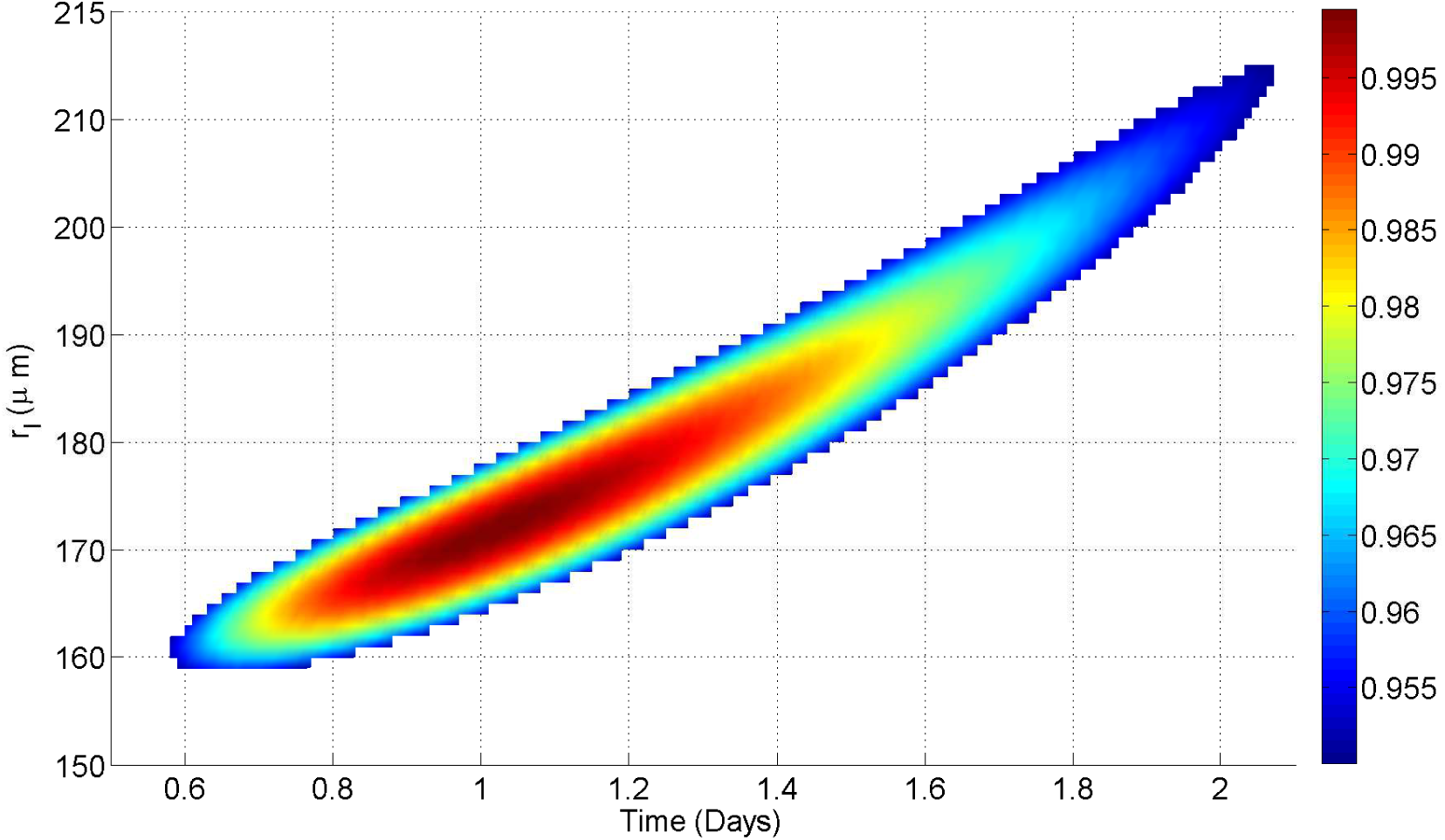
Best fit degeneracy for U-87 growth curve. While most values of *r_l_* / *t_d_* yield negative co-efficients of determination, there is a relatively narrow-band (shown in color) that produces a good fit to observed data (*R*^2^ > 0.95). In this case, values of *r_l_* between 160 - 215 *μ*m (8.56 - 15.46 ×10^−7^ m^3^ kg^−1^ s^−1^) can yield good fits, with these values yielding doubling times between 0.6 - 2.1 days. The range value is due to inherent degeneracy between diffusion limit and doubling time

For four cell lines, it was possible to use the OCR determination method outlined to obtain an estimate of OCR, circumventing degeneracy and testing the model further. With a known OCR, model growth curves could be directly contrasted to experimental data. This model validation was performed on growth curves for spheroids from a range of cell-lines, namely the HCT 116, LS 174T, MDA-MB-468 and SCC-25 cell lines. OCR was experimentally estimated using extracellular flux analysis combined with mass estimates derived from confocal microscopy and these estimates were then used to produce a growth curve, which was contrasted with measured spheroid growth curves for that cell line. From this, the best-fit cellular doubling time could be estimated and the agreement between experimental and theoretical curves quantified. For the cell lines analysed this way, agreement was high with mean co-efficient of determinations ranging from 0.9327 to 0.9958, suggesting the model is robust and describes the data well. This analysis was also attempted on the MDA-MB-231 and U-87 lines, but cell mass estimates of these lines were confounded by inability to accurately resolve individual cells, due either to high levels of mitotic cells (MDA-MB-231) or highly irregular cell shape (U-87). The SCC-25 line had considerable uncertainty in mass estimation (≥ 40%) which rendered its OCR uncertain and different from that in the pure curve fitting section.

Derived values for OCR lend themselves to estimations of cellular doubling time. Literature reports of doubling time have wide ranges, even for single cell lines, suggesting this may be heavily influenced by the conditions under which the cells are grown or the method used to estimate doubling time. For HCT 116, LS 174T, MDA-MB-468 and SCC-25, recent literature estimates of doubling time are 1.25 days, 1.33 days, 2 days and 2.1 days respectively [32–34]. This is in very good agreement with the estimated value for the LS 174T and MDA-MB-268 lines used in this work, but lower than estimated for the HCT 116 and SCC-25 lines. Interestingly, when SCC-25 was theoretically fit, the estimate for doubling time reduced to 2.85 days, closer to literature values than from the experimental case. There is also the possibility that cells in a spheroid grow differently to plated cells, and indeed there is some recent evidence for geometry specific metabolic differences in specific situations [35]. In theory, this could be tested by growing very small spheroids in their exponential phase of growth (*r_o_* << *r_c_*) and estimating the doubling time of the entire spheroid, however it is technically challenging to do so. It is also quite likely that the doubling time of cells is influenced by their growth conditions, which may render such comparisons void. In any case, the OCRs estimated from this method are in the same range as typical literature estimates. It was also possible to estimate HTC 116 OCR using two methods - this analysis yielded results within ≈ 1.7% of one another.

There are several potentially confounding factors in this work; one of which is the minimum oxygen tension required for mitosis, *p_m_*, taken to be 0.5 mmHg for all cell lines in this work [31]. It is possible however that hypoxic arrest limits differ between cell types. Higher values of would mean a decreased proliferating volume *V_p_* and consequently a decreased maximum volume. Conversely, lower values would act to increase *μ*. Whether this varies between cell lines is an open question. The effects of glucose are not modelled - this was because the spheroids were grown in a high glucose media (25mM or 4.5g/L) which suggests spheroids should be well supplied, and that the effects of low-glucose could be ignored without loss of generality. However, we can in principle use the model derived in this work to estimate what effect this might have; for low glucose environments, the literature estimate for minimal oxygen partial pressure for mitosis raises an order of magnitude to *p_m_* = 5 mmHg [31]. In this case, the proliferating volume *V_p_* would be markedly reduced. This situation is illustrated for a hypothetical spheroids in two different glucose concentrations in Fig B in supplementary material (**S1**), where it can be seen that even with an order of magnitude change in concentration, the effects are relatively small.

Another potentially confounding factor that may have considerable effect is the cellular density; in the results shown this was approximated to the density of water, but there is some evidence this can vary with cell cycle, and can be between 4% and 9% higher than the density of water [36]. If the higher estimates for density are used, this acts to increase the estimate of cellular mass and consequently reduces the estimated consumption rate as outlined in equation 5. For the fits shown in Fig 4 (a) - (c), this slightly improves the co-efficient of determination. In the case of SCC-25, this has the opposite effect, perhaps related to the suspect samples for this particular cell line. The fit data for higher density estimates is not shown for brevity.

The model makes an implicit assumption that the rate at which necrotic material is removed from anoxic core is sufficiently large that proliferation into the core region is not limited by space, and new cells are either pushed into available space in the core or bump existing cells into that void where they die due to insufficient oxygen. This inward migration of cells might be a consequence of several factors, but it is likely that high oxygen pressure around the spheroid or simply surface tension would act to confine the volume and push emerging cells into available central space. Were this not the case, one would expect spheroid radius to increase linearly and volume cubically without limit, which is not observed in practice, instead spheroids are seen to progress from linear growth to a plateau phase [3, 4]. This is also suggested by the model; initially the proliferating volume is greater than the anoxic core volume and so the spheroid experiences net growth, with an increasing anoxic core volume as cells die due to inadequate oxygen supply. Eventually, the anoxic volume is sufficiently large to absorb any new cells without net increase; equation 10 suggests that a plateau state is achieved when 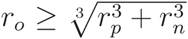 Re-arrangement of this identity shows this is the same as writing that plateau is achieved when the proliferating volume matches the anoxic core volume, as might be expected mechanistically. The exact kinetics of cell lysate removal from the core is not known, but imaging of sectioned spheroids appears to show an absence of cells in the centre, implying this material is indeed removed. When this condition is reached then even though the proliferating volume is constant and non-zero, the spheroids cannot increase in size. This situation is illustrated in the Fig C, supplementary material (**S1**).

While the model outlined here assumes lysis in the anoxic core occurs at a greater rate than cellular doubling time, it is also possible and worthwhile to explore the situation where the rate of lysis is such that the anoxic core is not totally removed and adds to the apparent spheroid volume. Supplementary figs D and E in supplementary material **S1** outline hypothetical situations where derived OCR and growth curves for cell lines used in this work were analysed under the assumptions of incomplete core removal. Specifically, a rate of half core removal and zero core removal. Using a similar approach outlined here, these resultant curves can be fit to the data, as outlined in supplementary Fig A in **S1**. However, under these assumptions, long term growth behaviour (illustrated in supplementary Fig E, supplementary material **S1**) deviates markedly from what is experimentally observed; under half core removal, spheroids would obtain unrealistically huge plateau radii, whilst spheroids without core removal would always increase exponentially and never plateau. These projections suggest that core removal is complete or close to complete for the spheroids analysed here.

It is also clear from the identity derived in equation 5 that cellular mass is needed to fully characterize OCR from extracellular flux analysis, and the range of masses estimated for cells in this work (2.12 - 3.54 ng) suggests cellular mass differs between cell lines, and can introduce substantial errors into OCR estimation if neglected. An example of this is seen in this work, where MDA-MB-468 cells produced higher values on the seahorse analyzer (Seahorse Bioscience, Massachusetts) than LS 174T cells, but due to the very small mass of the latter, the estimated OCR for LS 174T was in fact 14% higher than estimated OCR for the MDA-MB-468 cells.

The cell mass estimation in this work also has a number of potentially confounding factors. If cells are undergoing mitosis or apoptosis, these will tend to shew the volume estimates. To try and over-come this, both a membrane stain and nuclear stain were used so obviously unsuitable cells could be excluded from analysis, and a number of cells (22 ≤ *n* ≤ 36 ) were analyzed for each cell line. In other lines, cells were too close together to resolve (U-87) or all mitotic (MDA-MB-231) rendering the confocal volume approach of limited use. The volume reconstruction algorithm used here was relatively simple, and re-constructed volume estimates based on one micron slices. This might introduce errors for unevenly spaced cells. This could potentially be overcome with finer spacing, but this would act to increase bleed-through error and increase processing time. However, the majority of the derived mass estimates produced OCRs which matched the growth data well and mass validation with HeLa cells was encouraging.

The use of the model to estimate OCR change due to clinical compounds is also of interest. HCT 116 Spheroids treated with 50nM of Gemcitabine had a much lower OCR (mean 18.75 mmHg/s) than control HCT-116 spheroids (mean 27.43 mmHg / s) with OCR measured by the sectioning method [26]. Encouragingly, estimation of OCR for untreated HCT-116 spheroids by the sectioning method was in very close agreement (< 1.8%) to that determined in this work using the OCR technique outlined (mean 27.92 mmHg / s). The stark effect of Gemcitabine is also interesting; it is a known and potent radio-sensitizer [37–39], although the exact mechanism of action is still untested. This analysis might suggest that this drug reduces OCR, allowing greater oxygen diffusion and decreasing hypoxic regions, though more analysis would be required to test this hypothesis further. Even in vascular tissue, knowledge of the OCR and modulating influences on this have the potential to allow improved oxygen maps [40] and ultimately therapy delivery. The spheroids in this work were stained with the proliferation marker Ki-67, which shows all cell cycle phases except G0; this yields a binary distinction between quiescent and non-quiescent cells. For more detailed investigation of cell cycle state, more specific markers such as BrdU might be appropriate.

Spheroids in this work ranged in radius value from 182 *μ*m to 617 *μ*m, and OCR throughout growth was assumed to be constant for cell lines observed. This assumption is supported by analysis of DLD-1 colorectal spheroids which yielded an approximately constant oxygen consumption rate between 370 and 590 *μ*m [26], though analysis of other cell lines by different methods suggests that consumption rate can vary up to 50% in some lines whilst changing minimally in others [41]. This assumption is supported by a recent theoretical analysis [28] which examined hyperbolic Michaelis-Menten like oxygen consumption forms and found minimal variation between these forms and the simpler assumption of constant consumption rate. For spheroids in this analysis, a constant OCR assumption fits the data but if OCR variation is known for a given cell line it would be readily incorporated into the model. Although many types of spheroid culture methods exist, we used an ultra-low attachment format to grow spheroids for experimental simplicity and for volume measurements. The model presented here is relatively simple, and could be readily extended to incorporate other effects if the underlying parameters can be measured. There is also the possibility that oxygen dynamics differ between monolayers and spheroids, which would serve to confound the derived OCR data presented in table 2. This is an avenue worthy of further investigation, and it might be worthwhile to section stained spheroids to estimation their OCR by previously outlined methods [26] and contrast this with the OCR method outlined here. While there are numerous models in the literature for avascular spheroid growth [3,20-24], this model is novel as it specifically relates growth and growth limitation to oxygen status and availability.

## Conclusions

The model presented in this work yields projected spheroid growth curves from knowledge of oxygen consumption rate and cellular doubling time. Theoretical growth curves were found to match experimental data well for a range of cell lines with different characteristics, yielding the classic sigmoidal shape expected mechanistically without any a priori assumptions. This work also illustrates the importance of cellular mass is ascertaining OCR, and outlines a method for estimating this. Finally, a model was applied to infer the change in OCR due to clinical compounds, demonstrated with gemcitabine as a proof of principle.

## S1: Supplementary material

**Fig A**

**Growth curve arising from oxygen model** The assumption of oxygen mediated growth gives rise to the classic sigmoidal growth curve.

**Fig B**

**Growth curves as a function of glucose availability** Lower glucose levels give rise to a modified growth curve as *p_m_* increases when glucose isn’t available. Despite *p_m_* changing by up to a factor of 10 under such circumstances, the growth curves under conditions of both ample glucose and glucose deficiency are quite similar.

**Fig C**

**Relative radii of spheroid sections with growth** For a spheroid of initial radii 20 *μ*m and *a* = 7 × 10^−7^ m^3^ kg^−1^ s^−1^, the proliferating rim radius is the same as the spheroid radius. After *r* = *r_c_*, the proliferating radius falls relative to the anoxic radius *r_n_* which increases. At plateau, 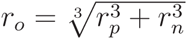 and no further increase in volume is observed.

**Fig D**

**Varying removal rate of anoxic core** It is possible to also model situations where only a fraction of the anoxic core is lysed relative to cellular doubling time. This can be illustrated on the longest term projections for which we have experimentally determined OCR data - (a) HCT 116 and (b) SCC-25. This was fit to the data and a best-fit estimate of cellular doubling time *t_d_* obtained, as illustrated.

**Fig E**

**Long term growth projections with different removal rates** Estimated projections for HCT 116 growth with time under different rate removal assumptions.

**Table A**

**Raw Seahorse oxygen consumption data for the HTC 116 cells used in this work.**

**Table B**

**Raw Seahorse oxygen consumption data for the LS 147T cells used in this work.**

**Table C**

**Raw Seahorse oxygen consumption data for the MDA-MB-468 cells used in this work.**

**Table D**

**Raw Seahorse oxygen consumption data for the SCC-25 cells used in this work.**

**Table E**

**Raw Seahorse oxygen consumption data for the MDA-MB-231 cells used in this work.**

**Table F**

**Raw Seahorse oxygen consumption data for the U-87 cells used in this work.**

## Acknowledgments

The authors gratefully extend their thanks to Prof. James Freyer for kindly allowing them use his V79 hamster cell spheroid data, and thank the reviewers for their suggestions on the kinetics of the necrotic core, and Dr. Daniel Warren of University of Oxford for his useful feedback.

